# Single cell transcriptional profiling of benign prostatic hyperplasia reveals a progenitor-like luminal epithelial cell state within an inflammatory microenvironment

**DOI:** 10.1101/2023.11.06.565375

**Authors:** Rei Unno, Jon Akutagawa, Hanbing Song, Keliana Hui, Yih-An Chen, Julia Pham, Heiko Yang, Franklin W. Huang, Thomas Chi

**Affiliations:** Division of Hematology and Oncology, Dept. of Medicine, University of California San Francisco; Helen Diller Family Comprehensive Cancer Center, University of California San Francisco; Bakar Computational Health Sciences Institute, University of California San Francisco; Institute for Human Genetics, University of California San Francisco; Dept. of Urology, University of California San Francisco; San Francisco Veterans Affairs Medical Center, San Francisco; Dept. of Nephro-urology, Nagoya City University Graduate School of Medical Sciences

## Abstract

Benign prostatic hyperplasia (BPH) is characterized by excessive cell proliferation and inflammation and affects most aging men. The development of new therapies for BPH requires a deeper understanding of the underlying pathophysiology and cellular components of BPH. Here, we characterize at single cell resolution the cellular states of BPH and identify cell populations enriched in BPH that contribute to cell proliferation and inflammation. Single-cell RNA-sequencing was performed on prostate tissue from 15 patients undergoing holmium laser enucleation of the prostate for treatment of BPH. Clustering and differential expression analysis on aligned single cell RNA-seq data was performed to annotate all cell types. Pseudotime, gene set enrichment, gene ontology, and ligand-receptor analyses were performed. 16,234 cells were analyzed and specific stromal, epithelial, and immune subgroups were found to be strongly associated with inflammation. A rare luminal subgroup was identified and pseudotime analysis indicated this luminal subgroup was more closely related to club and basal cells. Using a gene set derived from epithelial stem cells, we found that this luminal subgroup had a significantly higher stem cell signature score than all other epithelial subgroups, suggesting this subgroup is a luminal precursor state. Ligand-receptor interactions between stromal, epithelial, and immune cells were explored with CellPhoneDB. Unique interactions highlighting MIF, a pro-inflammatory cytokine that promotes epithelial cell growth and inflammation in the prostate, were found between fibroblasts and the progenitor luminal subgroup. This luminal subgroup also interacted with neutrophils and macrophages through MIF. Our single-cell profiling of BPH provides a roadmap for inflammation-linked cell subgroups and highlights a novel luminal progenitor subgroup interacting with other cell groups via MIF that may contribute to the inflammation and cell proliferation phenotype associated with BPH.

## Introduction

The prostate is a gland that produces seminal fluid in men and increases in size during aging in a disease process known as benign prostatic hyperplasia (BPH).^1–3^ An enlarged gland can obstruct the flow of urine and cause lower urinary tract symptoms (LUTS), which can significantly impact an individual’s quality of life.^2–4^ BPH negatively impacts approximately 50% of men aged 50 years, and its prevalence increases by about 10% each subsequent decade.^5^

BPH is thought to be due to abnormal glandular and stromal proliferation, primarily in the transition zone (TZ).^6,7^ This proliferation is associated with chronic inflammation and fibrosis, which are considered hallmarks of BPH.^8–10^ Macrophages and T cells have been implicated as the major inflammatory cell populations.^11,12^

Detailed cell states and interactions among these cell types are poorly understood as many previous characterizations have relied on unsorted bulk tissue samples and histological features.^13,14^ How inflammation drives epithelial and stromal proliferation or vice versa remains unclear. Without a better mechanistic understanding of BPH pathophysiology, better pharmacotherapeutic strategies cannot be developed. The efficacy of current BPH medications – alpha-blockers and 5-alpha-reductase inhibitors – is limited, and a large number of patients continue to rely on surgical BPH procedures for relief.^8,15^

Single-cell RNA-sequencing (scRNA-seq) enables the characterization of individual cell populations and prediction of their interactions at the transcriptional level.^16^ Recently, new prostate epithelial cell types, including hillock and club cells, have been detected in the normal and cancerous prostate using scRNA-seq.^17,18^ While most of these studies have focused on prostate cancer, BPH remains relatively unexplored.^19–21^ In this study, we used scRNA-seq to characterize the cell states and interactions associated with BPH. We identified cellular subgroups in the stromal, epithelial, and immune compartments strongly linked to inflammation. In particular, we found that a luminal epithelial subgroup had strong macrophage migration inhibiting factor (MIF)-driven interactions with stromal and immune cells. This epithelial subgroup and MIF could be key mediators of the inflammation and cell proliferation phenotype associated with BPH.

## Methods

### Experimental details

#### Sample collection

Fresh prostate tissue was obtained from fifteen patients undergoing holmium laser enucleation of the prostate (HoLEP) at the University of California San Francisco (UCSF) between January 2021 and January 2022. Patient demographics, comorbidities, and clinical data, including prostate size, PSA, and the presence or absence of 5-α reductase inhibitor usage, were obtained (Data Table 1). All specimens were confirmed pathologically to be non-cancerous in nature.

#### Study approval

The UCSF Institutional Review Board (IRB) committee approved the collection of the patient data included in this study. All relevant ethical regulations for work with human participants were compiled, and written informed consent was obtained.

#### Tissue collection and dissociation

Prostate tissue was enucleated using a Holmium laser^22^ and morcellated. Morcellated tissue was immediately delivered to the laboratory and further minced to ∼1-5mm^3^ pieces with surgical scissors in a petri dish with 10 mL RP-10 (RPMI + 10% FBS) on ice for up to 5 mins and washed with cold RP-10 (RPMI + 10% FBS). Each sample was centrifuged at 1200 rpm x5 mins at 4°C and resuspended in 10 mL warmed digestive media (HBSS + 1% HEPES) with 1,000 U/mL collagenase type IV (Worthington, Cat: LS004188), and rotated for 30 min at 37 °C. Samples were triturated by pipetting 10 times at the end of the incubation. Each sample was filtered through a 70-µm filter (Falcon, Cat: 352350), washed with RP-10, centrifuged at 1200 rpm x5 mins at 4°C, washed again with RP-10, and resuspended in RP-10. A hemocytometer was used to count the cells.^23,24^

#### Single-cell RNA sequencing

Sequencing was largely based on the Seq-Well S^3 protocol.^25,26^ One to two arrays were used per sample. Each array was loaded as previously described with approximately 110,000 barcoded mRNA capture beads (ChemGenes, Cat: MACOSKO-2011-10(V+)) and with 15,000– 20,000 cells. Arrays were sealed with functionalized polycarbonate membranes (Sterlitech, Cat: PCT00162X22100) and were incubated at 37 °C for 40 min.

After sealing, each array was incubated in lysis buffer (5 M guanidine thiocyanate, 1 mM EDTA, 0.5% sarkosyl, 1% BME). After detachment and removal of the top slides, arrays were rotated at 50 rpm for 20 min. Each array was washed with hybridization buffer (2 M NaCl, 4% PEG8000) and was then rocked in a hybridization buffer for 40 min. Beads from different arrays were collected separately. Each array was washed ten times with wash buffer (2 M NaCl, 3 mM MgCl_2_, 20 mM Tris-HCl pH 8.0, 4% PEG8000) and scraped ten times with a glass slide to collect beads into a conical tube.

For each array, beads were washed with Maxima RT buffer (ThermoFisher, Cat: EP0753) and resuspended in reverse transcription mastermix with Maxima RT buffer, PEG8000, Template Switch Oligo, dNTPs (NEB, Cat: N0447L), RNase inhibitor (Life Technologies, Cat: AM2696) and Maxima H Minus Reverse Transcriptase (ThermoFisher, Cat: EP0753) in water. Samples were rotated end-to-end, first at room temperature for 30 min and then at 52 °C overnight. Beads were washed once with TE-TW, once with TE-SDS, twice with TE-TW, and once with 10 mM Tris-HCl pH 8.0. They were treated with exonuclease I (NEB), rotating for 50 min at 37 °C. Beads were washed once with TE-SDS and twice with TE-TW. They were resuspended in 0.1 M NaOH and rotated for 5 min at room temperature. They were subsequently washed with TE-TW and TE. They were taken through second strand synthesis with Maxima RT buffer, PEG8000, dNTPs, dN-SMRT oligo, and Klenow Exo- (NEB, Cat: M0212L) in water. After rotating at 37 °C for 1 h, beads were washed twice with TE-TW, once with TE and once with water.

KAPA HiFi Hotstart Readymix PCR Kit (Kapa Biosystems, Cat: KK2602) and SMART PCR Primer (Supplementary Data 2) were used in whole transcriptome amplification (WTA). For each array, beads were distributed among 24 PCR reactions. Following WTA, three pools of eight reactions were made and were then purified using SPRI beads (Beckman Coulter), first at 0.6× and then at a 1× volumetric ratio.

For each sample, one pool was run on an HSD5000 tape (Agilent, Cat: 5067–5592). The concentration of DNA for each of the three pools was measured via the Qubit dsDNA HS Assay kit (ThermoFisher, Cat: Q33230). Libraries were prepared for each pool, using 1000 pg of DNA and the Nextera XT DNA Library Preparation Kit. They were dual-indexed with N700 and N500 oligonucleotides.

Library products were purified using SPRI beads, first at 0.6× and then at a 1× volumetric ratio. Libraries were then run on an HSD1000 tape (Agilent, Cat: 50675584) to determine the concentration between 400-800 bp. For each library, 3 nM dilutions were prepared. These dilutions were pooled for sequencing on a NovaSeq S4 flow cell.

The sequenced data were preprocessed and aligned using the dropseq_workflow on Terra (app.terra.bio). A digital gene expression matrix was generated for each sample, parsed, and analyzed following a customized pipeline.

### Quantification and statistical analysis

#### Alignment and QC

Sequencing results were received as paired FASTQ files. Each pair of FASTQ files was aligned against GRCh38 reference genome using a WDL-based Dropseq workflow (https://cumulus.readthedocs.io/en/latest/drop_seq.html) on Terra. All default parameters were used in the alignment except the parameter –dropseq_tools_force_cells set as 2,000 to ensure maximum cell capture. The alignment results include aligned and corrected bam files, gene expression matrices in .txt.gz format, and txt files of alignment reports such as the list of barcodes, the number of reads, and alignment summaries. For each sample, the median and average number of genes captured per barcode were 419 and 628. The median and average number of UMIs captured per barcode were 818 and 1,511. The average percentage of mitochondrial content per barcode was 23.5%. Each gene expression matrix was then processed through decontX^27^ for ambient RNA-decontamination. Barcodes with less than 300 genes, 500 UMIs, or more than 20% of mitochondrial content were removed from downstream analysis. Eventually, a total number of 16,234 cells were analyzed with an average of 951 genes, 1,598 UMIs and 6.4% of mitochondrial content.

#### Clustering analysis

Single-cell cluster analysis was performed with custom scripts utilizing the Seurat package (v4.3.0) in R (v4.2.0) and Scanpy package (v1.7.2) in Python (v3.6.8). Data from the samples were merged, as integration of the data did not discernibly alter cell clustering analysis. Individual cell UMI-collapsed read count matrices were loaded for analysis. We removed 2,850 genes that were detected in 3 or less cells and all mitochondrial genes from downstream analyses. Highly variable genes with a minimum and maximum gene expression of >=0.0125 and <= 3, respectively and a minimum dispersion of genes = 0.5 were selected. Gene expression of all cells was normalized by multiplying by a scale factor of 10,000 and subsequently log-transformed. Using 50 calculated principal components, we identified 26 clusters. The Leiden algorithm was used for clustering. Marker genes were defined by a Wilcoxon pairwise differential expression analysis in Scanpy. Feature and violin plots were used to visualize gene expression.

#### Differential gene analysis

Differential gene analysis was performed using the rank_genes_group module in Scanpy, which performs a Wilcoxon rank sum test between groups.

#### Pseudotime analysis

To assess the epithelial cell states regarding their orders in the differentiation trajectory, we conducted pseudotime analysis on all epithelial cells in the HoLEP scRNA-seq dataset using Monocle3 (https://cole-trapnell-lab.github.io/monocle3/). Based on previously established studies, we selected basal cells as the starting point and computed the pseudotime trajectory. Specific gene expression levels along the trajectory were also visualized.

#### Gene ontology and gene set enrichment analysis

The package GSEAPY (v1.0.4) was used to predict differences in pathway enrichment between subgroups or phenotypes. Gene sets from MSigDB served as references for these analyses.

#### Ligand receptor analysis

The package CellPhoneDB (v3.1.0) was used to predict significant ligand-receptor interactions between different cell types. Cell subgroups were downsampled to 200 cells prior to analysis. Significant mean and cell communication significance (p < 0.05) was calculated for cell subgroups and patients.

## Results

### Single cell transcriptomic landscape of benign prostate hyperplasia

Prostate tissues from BPH patients undergoing Holmium laser enucleation of the prostate (HoLEP) were dissociated into single cells and then captured and sequenced using the Seq-well single-cell RNA-sequencing platform.^25,26^ After ambient RNA decontamination and removal of low-quality cells, we captured 16,234 cells from 15 different patients (Data Table 1) for analysis (Figure 1a). Using a graphical clustering method, 14 clusters of distinctive transcriptomic profiles were initially identified using a semi-supervised annotation pipeline (Figure 1b). Differentially expressed genes (DEGs) in each cluster were computed, and canonical cell-type-specific markers were used to guide the annotation of epithelial cells, endothelial cells, stromal cells, and immune cells (Figure 1d).

**Fig. 1.**
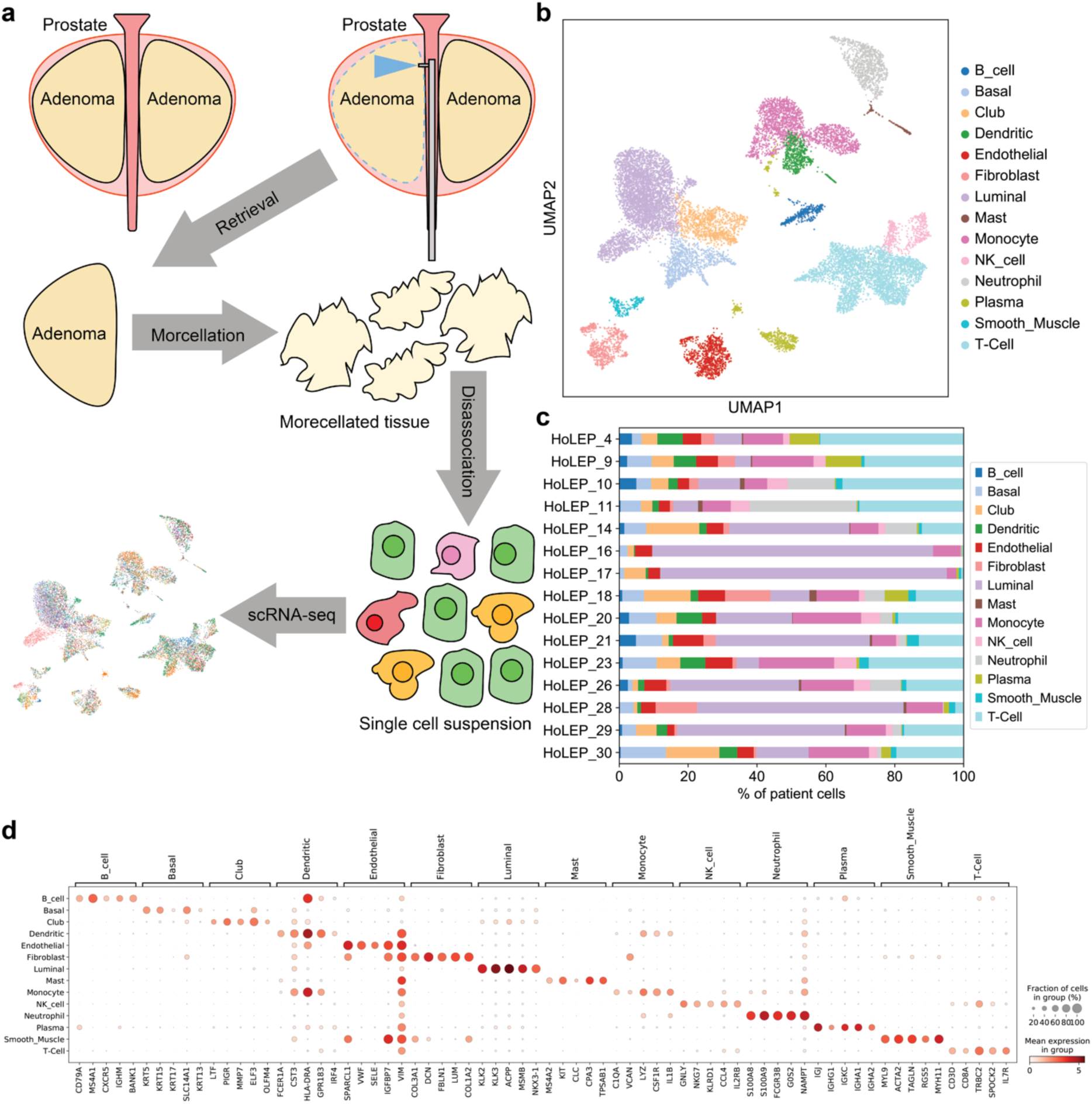
Identification of 14 cell groups in BPH samples via single cell RNA-sequencing. (a) Schematic of single cell RNA-sequencing of BPH cells obtained via HoLEP. (b) Uniform manifold approximation and projection (UMAP) of BPH transcriptomes show 14 distinct cell clusters. (c) Cell type composition of 15 BPH samples. (d) Dot plot of expression levels of known marker genes in each cell type.

Epithelial cells were the most abundant population and could be separated into three different subtypes, including basal, club, and luminal epithelial cells. Basal cells were enriched in basal markers such as *KRT5*, *KRT15*, and *TP63*, and club cells were enriched with previously established club cell markers and intermediate prostate epithelial cell markers such as *PIGR*, *MMP7*, *LTF*, and *SCGB3A1*. Luminal cells, identified by expression of *KLK3*, *KLK2*, and *ACPP*, were the largest epithelial population detected (n = 4,302 cells).

As with epithelial cells, immune cell analyses also revealed several subtypes. T cells that expressed *CD8A* and *CCL5* were the most abundant immune cell type present (n = 3,370 cells), followed by monocyte cells (*C1QA* and *IL1B*, n = 1,952), neutrophils (*S100A8* and *S100A9,* n = 892), dendritic cells (*FCER1A*, n = 585 cells) and then plasma cells (*IGJ* and *IGHA1*, n = 490 cells). Stromal cells consisted of fibroblasts (*FBLN1* and *DCN*) and smooth muscle cells (*ACTA2* and *MYLA9*). Most cell types received contributions from multiple patients and allowed a cell type composition comparison (Figure 1c). No significant associations were found between broad cell type composition and clinical features (ancestry, patient age, prostate size) in this cohort.

### Distinct fibroblast groups associated with inflammation pathways

Among all cell types, fibroblasts were the cell type with the strongest enrichment of a BPH signature gene set^14^ (Figure 2a). To further investigate this population, we used unsupervised clustering to define four fibroblast groups and two smooth muscle groups (Figure 2b). Canonical fibroblast markers – *LUM*, *VCAN*, and *BMP5* – were expressed in all fibroblast clusters. Smooth muscle populations were enriched in genes such as *ACTA2*, *MYH11*, and *MYL9*, which are all smooth muscle specific markers^17^. Based on the DEG analysis, we characterized the four fibroblast groups by their marker genes (Figure 2c, Supplemental Data 2). We found that the fibroblasts expressing *C7* were most enriched in genes found in the BPH signature gene set (Figure 2d,e). *C7*, which encodes for complement component C7, has been previously reported as a marker for interstitial fibroblasts in BPH samples.^20^ The *CCL2*-expressing fibroblasts were also highly enriched in BPH-associated genes in this cohort.

**Fig. 2.**
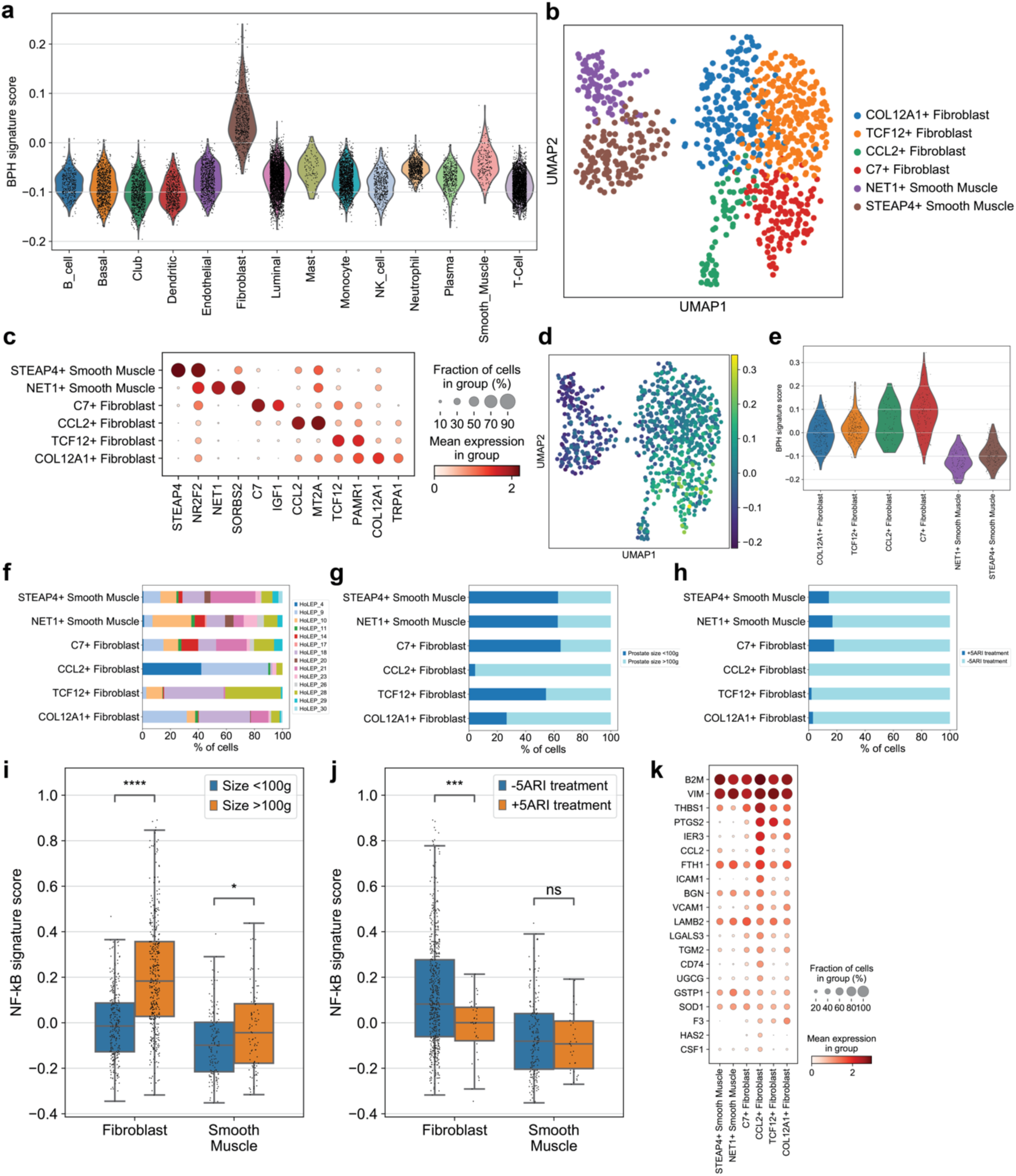
Single cell profiling of stromal cell compartment in BPH patients. (a) Violin plot of BPH signature score, derived from Middleton et al., across all cell groups. (b) UMAP clustering of stromal cells. (c) Dot plot of selected differentially expressed genes across the 6 stromal subtypes. (d) Feature plot of stromal cells, showing BPH-associated gene expression of each cell. (e) Violin plot of BPH signature score across the 6 stromal subtypes. (g) Cell number ratios of the 6 stromal groups, calculated by prostate size. (h) Cell number ratios of the 6 stromal groups, calculated by 5-ARI treatment status. (i) NF-κB pathway gene signature score of fibroblast and smooth muscle cells by prostate size. (j) NF-κB pathway gene signature core of fibroblast and smooth muscle cells by 5-ARI treatment status. (k) Dot plot of NF-κB pathway associated gene expression in the 6 stromal subtypes.

The C7+ and COL12A1+ fibroblast clusters consisted of cells from 12 patients, while the CCL2+ fibroblast cluster consisted of cells mostly from patients 4 and 9 and the TCF12+ fibroblast cluster consisted of cells mostly from patients 10, 18, and 28 (Figure 2f, Supplemental Data 2a). We observed that BPH patients with prostates larger than 100g harbored more fibroblasts expressing *CCL2* and *COL12A1* (Figure 2g) than those less than 100g (p < 0.05, Mann-Whitney U test). We also observed smaller populations of fibroblasts in patients treated with 5-alpha reductase inhibitors (5-ARIs) (Figure 2h). The majority of fibroblasts from 5-ARI-treated patients lacked CCL2+ and COL12A1+ fibroblasts, suggesting these two groups of fibroblasts may be localized in regions that undergo atrophy following 5-ARI treatment. Expression of genes associated with the inflammatory TNF-alpha/NF-κB pathway was higher in fibroblasts and smooth muscle cells from patients with larger prostates (Figure 2i) (p < 1e-5, Mann-Whitney U test). Fibroblasts from patients treated with 5-ARIs had lower expression of inflammation-associated genes (Figure 2j) (p < 0.001, Mann-Whitney U test). Specifically, CCL2+ fibroblasts had the highest expression of NF-κB pathway genes, while C7+ fibroblasts had the lowest (Figure 2k, Supplemental Data 3e). Taken together, these data suggest that fibroblasts in prostates with BPH, particularly those expressing *CCL2*, exhibit an inflammatory phenotype associated with the disease.

### A progenitor luminal epithelial cell subpopulation

As epithelial-to-stromal crosstalk is a known feature in BPH, we hypothesized that a fibroblast cell state in BPH may harbor specific interactions with surrounding epithelial cells to promote inflammation or cell proliferation.^28^ To investigate this possibility, we performed a subgrouping analysis of epithelial cells and identified subgroups in basal cells, club cells, and four subgroups of luminal epithelial cells (Figure 3a). The basal cell subgroup strongly expressed basal markers *KRT5* and *KRT15* and club cells expressed *PIGR*, *LTF*, and *MMP7*, consistent with previous scRNA-seq studies of human prostates.^17^ All four luminal subgroups shared high expression of canonical luminal epithelial markers, including *KLK3*, *KLK2*, *ACPP*, and *NKX3-1* (Figure 3c) and genes associated with the androgen receptor pathway. These four luminal subgroups included two larger subgroups (n=2,026 and 1,495) that were largely overlapping in gene expression profiles, a third subgroup (n=595) found in only one patient, and a much smaller fourth subgroup (n=186) that had contributions from multiple patients and was characterized by higher expression of *KLK4*, *HOXB13*, and *SORD* (Figure 3c, Supplemental Data 4). Significantly upregulated genes in luminal subgroup 4 compared to the first two luminal subgroups also featured many ribosomal genes (Supplemental Data 4), including *RPS15*, *RPL27A*, *RPS2*, *RPL7A*, and *RPLP0*. The top significantly enriched gene set from a gene ontology (GO) analysis of this subgroup was a ribosomal signature (Supplemental Data 5). This subgroup also highly expressed *FTH1* and *FTL*, genes that are both associated with a protective role in ferroptosis-mediated proliferation.^29^

**Fig. 3.**
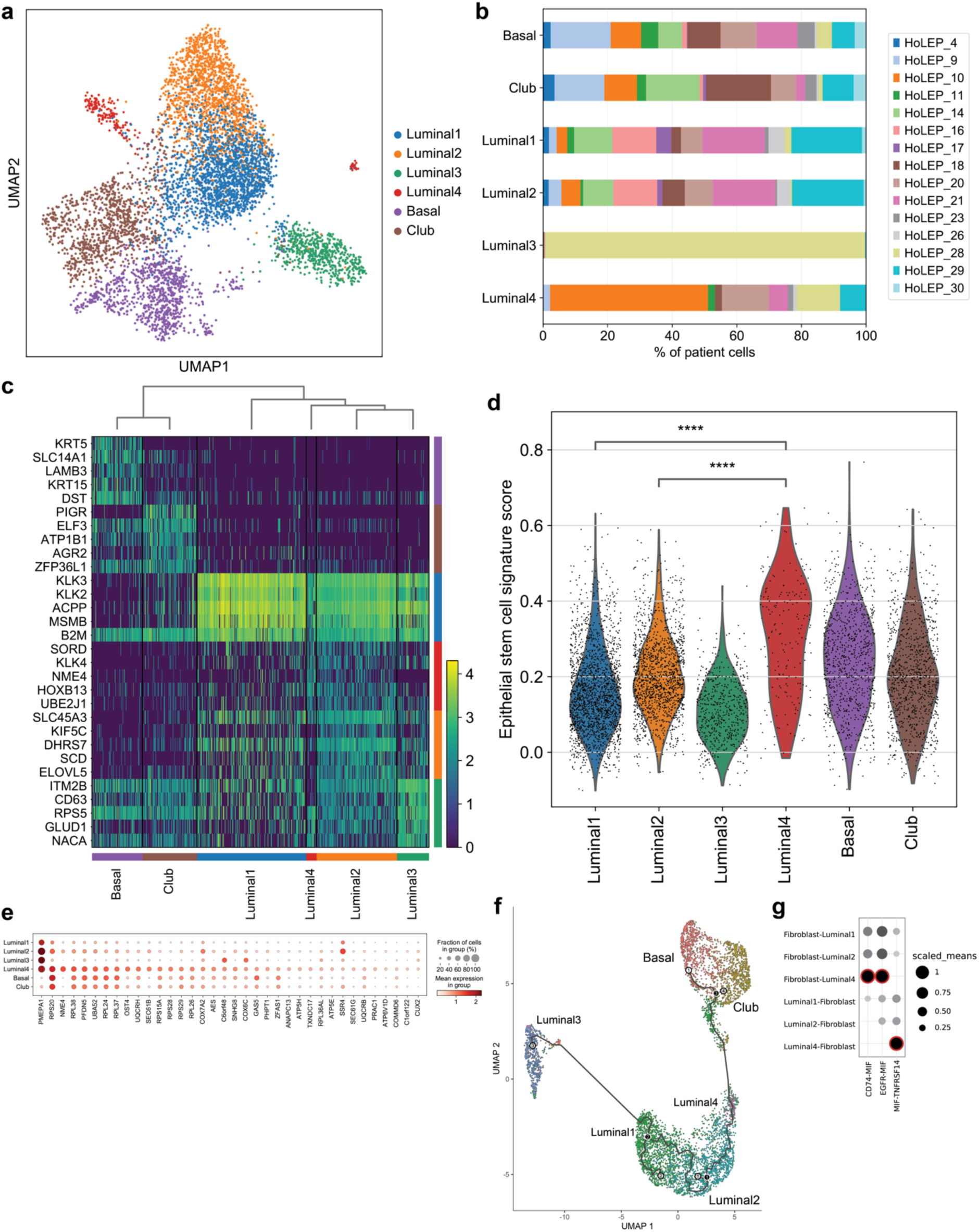
A progenitor luminal subgroup interacts with fibroblasts through MIF. (a) UMAP clusters epithelial cells into 6 subgroups. (b) Cell contributions from each sample to the epithelial subgroups. (c) Heatmap of differentially expressed genes in epithelial cells. (d) Violin plot of epithelial stem cell-associated gene expression scores in the 6 epithelial subgroups. (e) Dot plot of stem cell-associated gene expression in the 6 epithelial subgroups. (f) Trajectory analysis of the 4 luminal subgroups. (g) Interaction analysis between fibroblasts and luminal cells. Larger dot sizes represent higher means of average expression level of the two interacting molecules. Red highlights represent a significant interaction, with an adjusted p-value <0.05.

Using a geneset derived from epithelial stem cells^30^, we also found that the luminal 4 subgroup had a significantly higher stem cell signature score compared to all other epithelial subgroups (p < 0.00001, Mann-Whitney U test), including the basal and club cell groups (Figure 3d). *PRAC1*, a known prostate stem cell marker, is also uniquely expressed in this luminal subgroup (Figure 3e). To further characterize the 4 luminal subgroups, we performed a pseudotime analysis. Using basal cells as the starting point to construct the pseudotime trajectory, we found that the luminal 4 subgroup gave rise to the other luminal subgroups, suggesting that it may represent a luminal progenitor cell state (Figure 3f). Cell cycle regression analysis found no cell cycle differences between the progenitor luminal cells and other luminal subgroups. Genes highly expressed in the luminal 4 subgroup were used to create a geneset signature and were used to detect this luminal progenitor subgroup in an orthogonal prostate cancer dataset (Supplemental Data 6). It expressed similar genes to a tumor cell subset, suggesting that this progenitor subgroup may also be a tumor precursor.

To investigate the interaction of these epithelial cells with the stromal population and fibroblasts specifically, we performed ligand-receptor interaction analysis between the two stromal populations (fibroblasts and smooth muscle cells) and the three luminal subgroups (subgroup 1,2 and 4) using CellPhoneDB (Figure 3g). Dihydrotestosterone (DHT) to androgen receptor (AR) was predicted to be a significant interaction between the fibroblast subgroup and all luminal subgroups, consistent with previous studies.^31,32^ We detected four significantly increased interactions between the luminal 4 subgroup and fibroblasts (Supplemental Table 5). 3 of these 4 interactions featured macrophage migration inhibitory factor (MIF).

MIF is a macrophage-associated proinflammatory factor that is overexpressed in prostate cancers.^33^ *MIF* was found to be highly expressed in only the luminal 4 group, compared to the other luminal groups (Supplemental Data 5) and the complex formed with its main receptor, CD74, was predicted to be a significant interaction between the luminal 4 subgroup and fibroblasts. Epidermal growth factor receptor (EGFR) to MIF was also predicted to be a significant interaction between fibroblasts and the progenitor luminal subgroup alone but not the other two luminal subgroups. Between the fibroblasts and progenitor luminal cells, MIF was predicted to interact with TNFRSF14, which is a cell surface receptor and member of the tumor necrosis factor receptor superfamily that drives inflammation and immune response.^34^ Consistent with our hypothesis, we identified a luminal subgroup that may act as a progenitor cell state with specific interactions strongly associated with inflammation with surrounding fibroblasts and smooth muscle cells.

### Immune cell analysis reveals 3 distinct inflammation-related macrophage subtypes

In our immune cell analysis, we identified 13 distinct groups of immune cells with distinct transcriptomic profiles via unsupervised clustering (Figure 4a). Lymphoid (IL7R+) and myeloid lineages (CD14+) were predominant and T cells comprised the largest portion of the lymphoid groups, particularly the CD4+ (n=1,177) and CD8+ (n=1,728) subgroups. Using marker genes from Tuong et al.,^35^ we identified CD4 tissue resident memory (TRM) cells (n=465), CD16+ natural killer cells (n=276), CD16-natural killer cells (n=219), and B cells (n=363). We also identified plasmacytoid dendritic cells (n=51), neutrophils (n=892), mast cells (n=115), and CD86+ dendritic cells (n=534). All subgroups had contributions from the majority of patients (Figure 4b), except for neutrophils, with contributions from 9 patients.

**Fig. 4.**
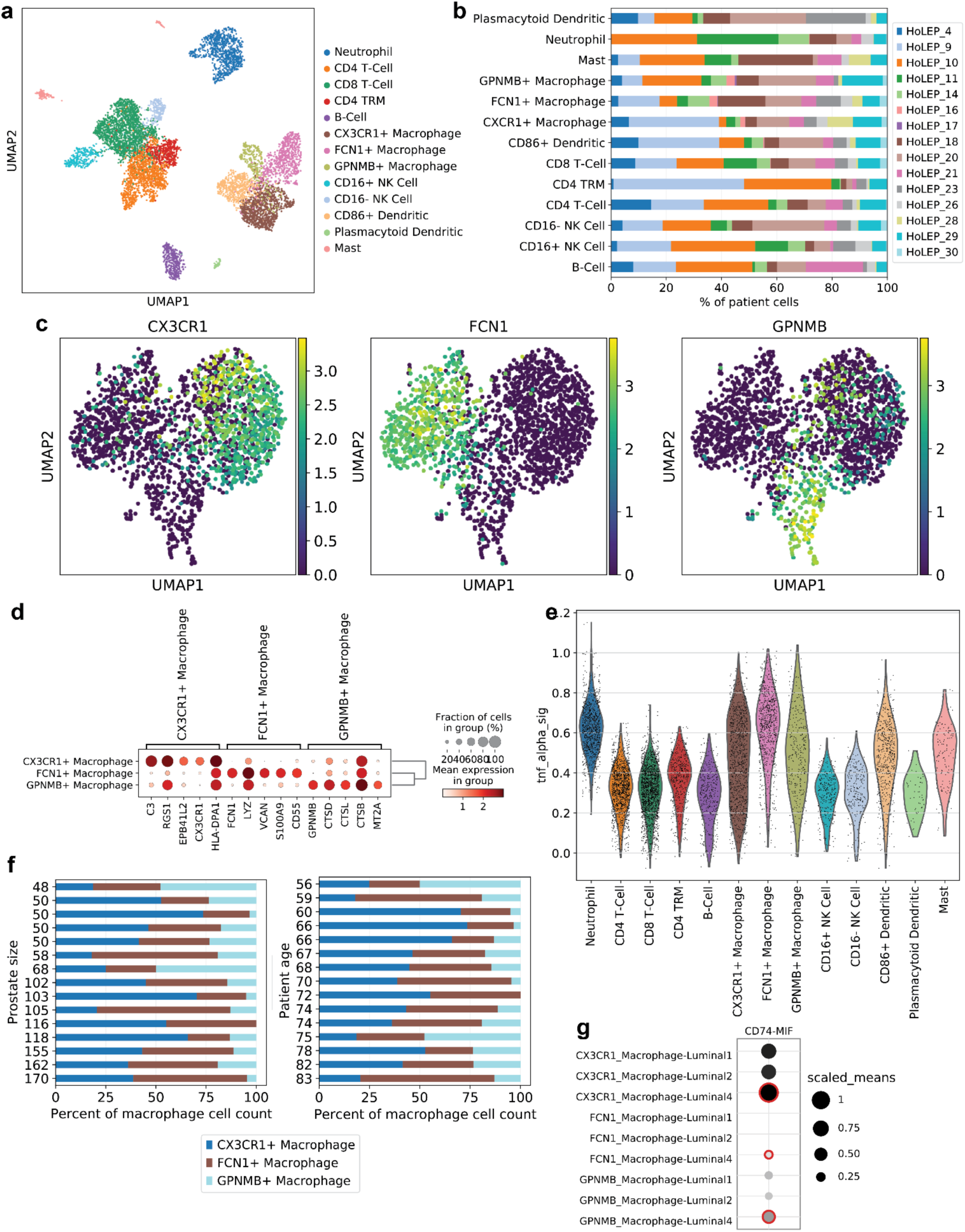
MIF-related macrophages and neutrophils are linked to inflammation. (a) UMAP shows clustering of immune subtypes. (b) Patient contribution to each immune cell subtype. (c) Feature plots of *CX3CR1*, *FCN1*, and *GPNMB* expression in macrophages. (d) Dot plot of marker gene expression in the three macrophage subgroups. (e) Violin plot of TNF-alpha pathway gene expression across all immune cell types. (f) Macrophage subtype composition ordered by prostate size (cc) or patient age. (g) Interaction analysis of luminal subgroups and macrophages. Larger dot sizes represent higher means of average expression level of the two interacting molecules. Red highlights represent a significant interaction, with an adjusted p-value <0.05.

Macrophages were the second most abundant group of immune cells. Three separate macrophage groups were identified using differential gene expression analysis (Figure 4c,d). One group strongly expressed *CX3CR1*^36^ and *C3*^37^ (n=937), which are both inflammation regulators in macrophages. Another macrophage group expressed *FCN1*, *VCAN*, and *S100A9* (n=716), which are also previously noted as inflammatory markers.^38^ The third group expressed *GPNMB*^39^ and *MT2A*^35^ (n=299), which have been associated with changes in prostate tumor microenvironments. The three macrophage groups strongly expressed genes associated with the M1 inflammatory phenotype. M2 anti-inflammatory macrophages were largely absent from these prostatic samples (Supplemental Data 7). Using genes from the TNF-alpha and NF-κB pathway, we found a strong inflammation signature in all three macrophage groups, neutrophils, CD86+ dendritic cells, and mast cells (Figure 4e). Gene set enrichment analysis of genes differentially expressed in the three macrophage populations revealed FCN1+ and CX3CR1+ macrophages to be enriched in inflammation-associated genes (Supplemental Data 8) - these two groups were often the largest share of macrophages regardless of prostate size or patient age, further supporting the inflammation phenotype associated with BPH (Figure 4f). Cell communication analysis between the epithelial and immune cells predicted interactions between macrophages and luminal subgroup pairs. The CD74-to-MIF interaction was predicted with all three macrophage groups and the progenitor luminal group, with the CX3CR1-Luminal 4 interaction being the strongest (Figure 4g). We also found additional unique interactions predicted to occur between MIF expressed by the progenitor luminal subgroup and two TNF-alpha receptors, TNFRSF14 and TNFRSF10D, expressed by the macrophages. The neutrophils and progenitor luminal subgroup also were predicted to have similar significant interactions featuring MIF, CD74, and TNFRSF14 (Supplemental Data 8).

## Discussion

Our study is the first analysis of BPH cells collected by HoLEP. With HoLEP becoming the standard for surgical treatment of BPH,^40^ samples generated by the procedure can reveal biological insights about the origins of diseases in the prostate. Here we used single cell transcriptomic profiling to characterize samples extracted through HoLEP and found multiple stromal, epithelial, and immune subgroups that were linked to inflammation. CCL2 is a chemoattractant that attracts both myeloid^41^ and lymphoid cells under different contexts^42^ and is involved in psoriasis,^43^ atherosclerosis,^44^ and other inflammation-related diseases.^41^ Interestingly, CCL2 is not typically expressed in normal prostate fibroblasts, but is a biomarker for prostate tumor microenvironments.^45^ Increased CCL2 has been associated with the proliferation of prostate stromal cells^46^ and is highly expressed in BPH fibroblasts. It has also been implicated as a potential factor in prostate cancer cell migration and infiltration.^47^ CCL2+ fibroblasts in our study had the highest expression of NF-κB pathway genes compared to all other stromal cells and may be associated with the sustained cell recruitment and inflammation found in BPH stromal cell layers. The FCN1+ and CX3CR1+ macrophage subgroups in this cohort also have strong inflammatory profiles and are driven by known immunomodulatory genes associated with classical M1 macrophages. In particular, CX3CR1 has been reported as a potential drug target for inflammatory diseases like psoriasis^48^ and atherosclerosis.^49^

Ligand-receptor interaction analysis predicted unique interactions involving MIF between the progenitor luminal subgroup and fibroblast, macrophage, and neutrophil subgroups. MIF is a pro-inflammatory cytokine that plays a regulatory role in both the innate and adaptive immune responses.^50^ Some studies have shown that MIF can promote tumor growth and other diseases.^51,52^ With regard to the prostate, luminal epithelial cells express MIF and MIF expression is significantly higher in BPH epithelium compared to normal epithelium.^53^ Modulating MIF overexpression may also lead to better clinical outcomes for BPH patients.^54^ MIF can also affect signaling cascades through its binding to CD74. CD74 is heavily involved with cell survival and proliferation^55^ and this complex activates the mitogen-activated protein kinase (MAPK) and phosphoinositide 3-kinase (PI3K) pathways associated with proliferation and angiogenesis. ^55^ MIF signaling can also increase the expression of the inflammatory cytokines IL-6 and IL-8^56^ and CD74 directly interacts with CXC-chemokine receptor 2 (CXCR2) to form a receptor complex for IL-8 that can potentially result in an inflammatory feedback loop.^57^ Our current study predicted CD74-MIF as a unique interaction between multiple cell types and the progenitor luminal subgroup, suggesting that this progenitor luminal subgroup may contribute to the BPH phenotype.

The progenitor luminal subgroup may represent a novel epithelial cell state or subtype associated with uncontrolled growth and proliferation. Compared to the other luminal subgroups, these cells highly express KLK4, which has been characterized as a potential biomarker for prostate, breast, and ovarian cancers. ^58–60^ The regulation of KLK4 by androgens^61^ and its function as a proliferative factor in the development of prostate cancer^62^ suggest a possible role for this KLK4-high luminal subgroup in the promotion of prostatic enlargement. In addition, gene ontology analysis revealed a significant enrichment of established ribosomal gene sets in these KLK4-high cells. As ribosomal biogenesis is a crucial biological process closely tied to cell growth and proliferation,^63^ this luminal subgroup could be associated with the development of prostatic hyperplasia.

Through a pseudotime analysis, we also found the gene expression dynamics of this luminal cell group were more similar to the club and basal cells than the other luminal subtypes. While the size of this cell group is relatively small, we hypothesize this cell group represents a luminal progenitor state. The majority of prostate stem cells are found in the basal cell group,^64^ so these luminal cells may constitute a progenitor subtype that contributes to the inflammation phenotype and proliferation of the stromal and epithelial cells.

As BPH is a disease of inflammation,^8–11^ we detected three macrophage subgroups in immune cell populations associated with inflammation. All macrophage groups were associated with the M1 inflammatory phenotype and had a strong inflammation signature. The 3 macrophage groups, particularly the CX3CR1+ macrophages were shown to have a strong interaction with the progenitor luminal cells, potentially demonstrating how these luminal cells drive inflammation through macrophage activation.

Currently, alpha-adrenergic blockers and 5-alpha reductase inhibitors are used to treat BPH.^65^ While alpha-blockers can minimize lower urinary tract symptoms by inhibiting smooth muscle contraction, 5-ARIs reduce prostate volume by decreasing dihydrotestosterone production. Although 5-ARI has been shown to improve clinical outcomes successfully, not all patients respond and some develop resistance.^66^ Recent data indicate that atrophy of prostate luminal cells caused by 5-ARI correlated with reduced AR signaling and increased NF-κB signaling. 5-ARI also induced a luminal-to-club cell transition.^67^ In our study, while the inflammatory TNF-alpha/NF-κB pathway was higher in fibroblasts from patients with larger prostates, fibroblasts from patients treated with 5-ARIs had lower expression of these inflammation-associated genes. Across all fibroblast groups, fibroblasts from patients that were previously treated with 5-ARIs were lower in abundance than untreated patients. AR signaling was increased in patients with large prostate volumes and decreased in those with 5-ARI treatment.

Our results suggest the MIF-CD74 interaction as a potential target in BPH patients who do not respond to 5-ARIs, particularly ones that exhibit high inflammation, and may benefit from next-generation MIF targeting drug development. Targeted MIF inhibitors, such as CPSI-1306,^68^ benzoxazole-2-thione,^69^ and *N*-acetyl-*p*-benzoquinone imine,^70^ have demonstrated the ability to disrupt the binding of MIF to CD74. These inhibitors have shown some promise in clinical trials for cancer and autoimmune disease^53,54,56^ and may be potential therapeutic options for BPH. Due to the increasing popularity of HoLEP as a treatment method for BPH, our study demonstrates that analyses on cells collected from this procedure can further contribute to our understanding of BPH.

## Conclusions

This study provides a single-cell transcriptomic profile of BPH and highlights stromal and immune cells that are strongly associated with inflammation. A novel luminal epithelial subgroup was also identified as a potential driver for this inflammation phenotype through the expression of *MIF*. MIF interacts with cell surface receptor proteins on stromal and immune cells and may contribute to recruitment and proliferation of these cells. While further studies are needed to confirm these luminal progenitor cells and MIF as new biomarkers and targets for BPH, our results show that single cell analyses can identify unique cell types and interactions associated with BPH.

## Supplemental Data

*Data Table 1 - Patient data*

*Data Table 2 - Gene signatures for 14 clusters*

*Data Table 3 - Gene signatures for 6 stromal subgroups*

*Data Table 4 - Gene signatures for 3 macrophage subgroups*

*Data Table 5 - Significant interactions from CellPhoneDB*

**Supplemental Data 1.**
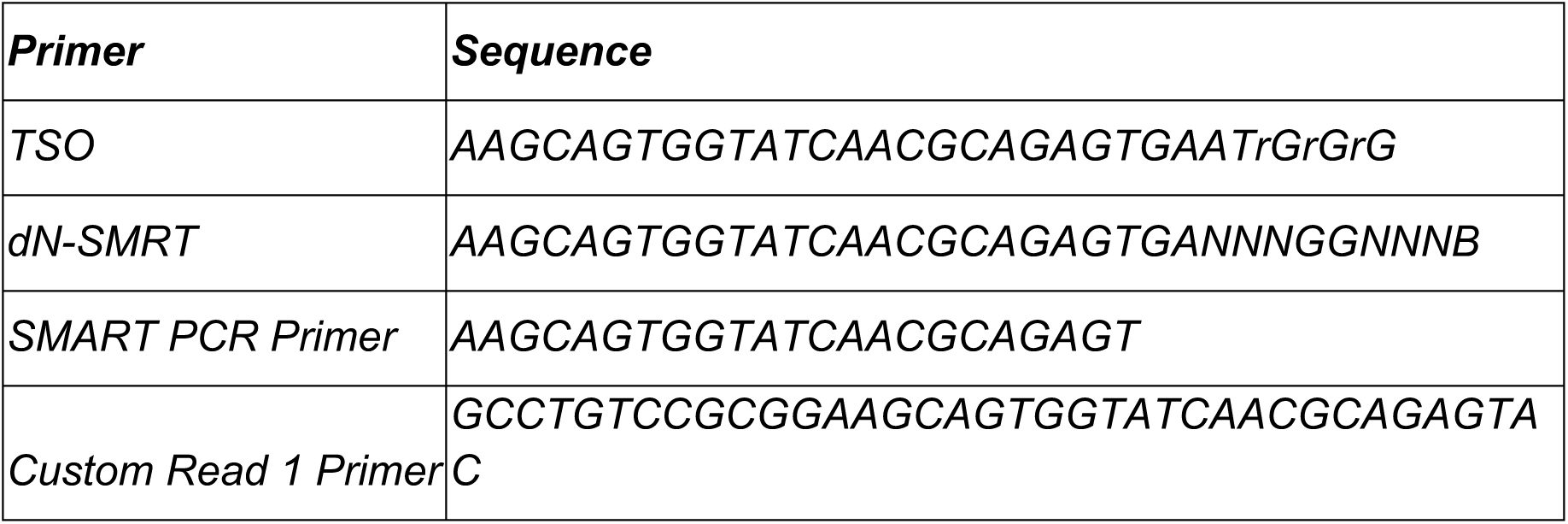
SMART PCR Primer. Primer sequences used for whole transcriptome amplification.

**Supplemental Data 2.**
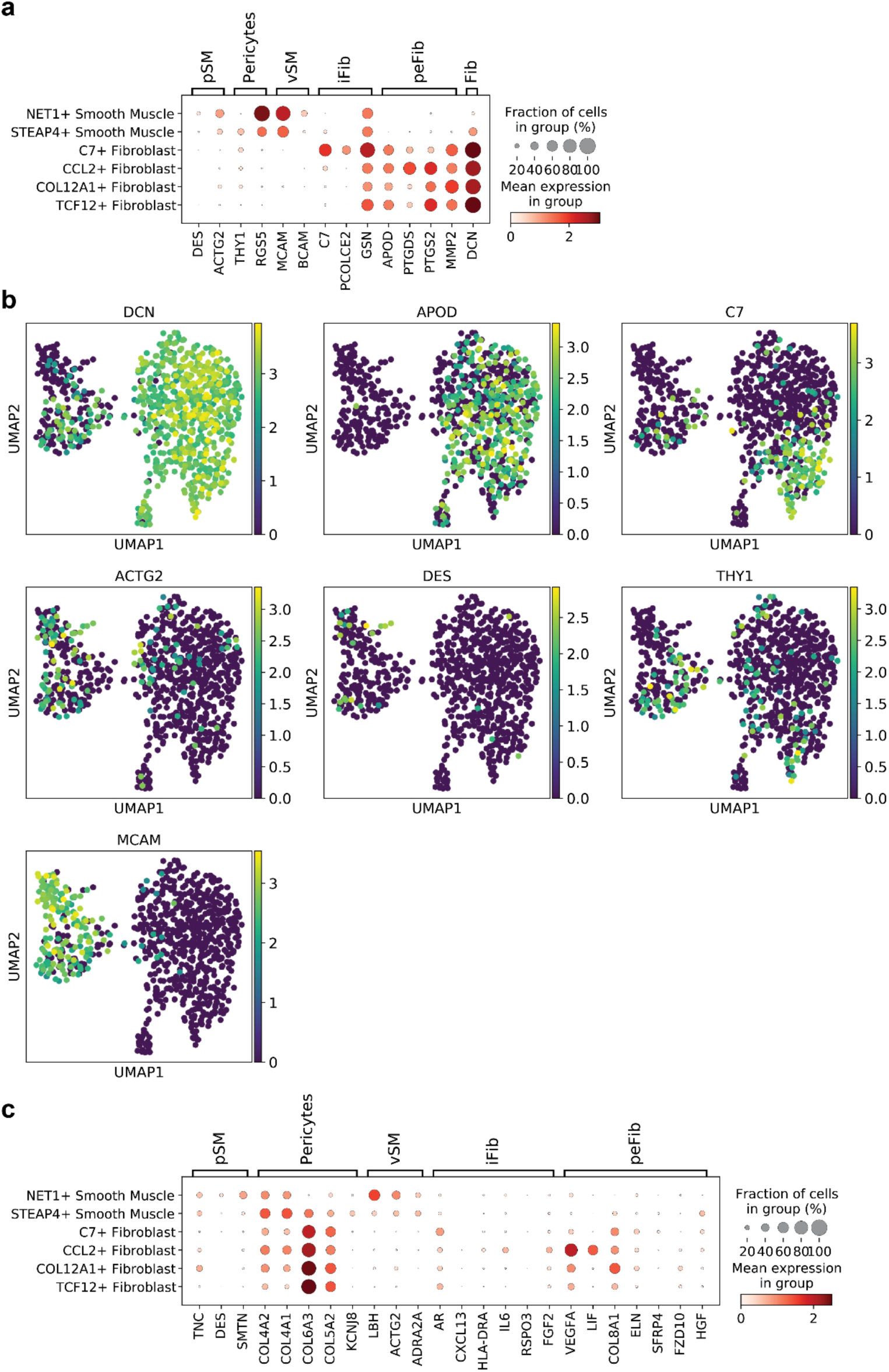
Stromal cell marker comparison. (a) Dot plot of stromal marker genes highly expressed in normal prostate samples from Joseph et al., 2021. (b) Feature plots of previously described marker genes. (c) Dot plot of stromal marker genes highly expressed in BPH samples from Joseph et al., 2021.

**Supplemental Data 3.**
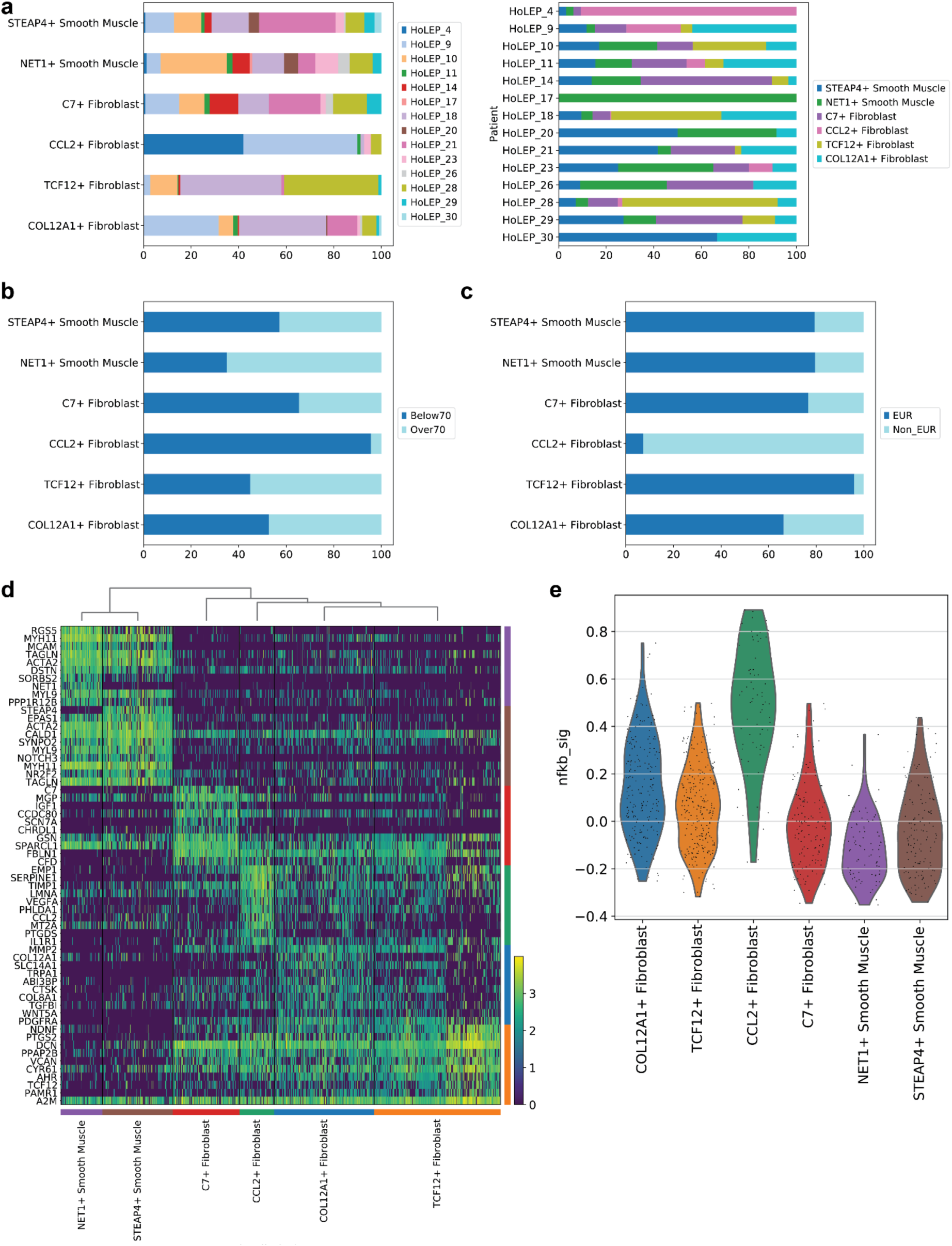
Stromal cell characterization. (a) Cell contributions from each patient. (b) Proportion of stromal cells in patients under or over age 70. (c) Proportion of stromal cells in patients based on ancestry. (d) Heatmap of differentially expressed genes from each stromal cell group. (e) Signature score derived from NF-κB pathway associated genes of each stromal cell group.

**Supplemental Data 4.**
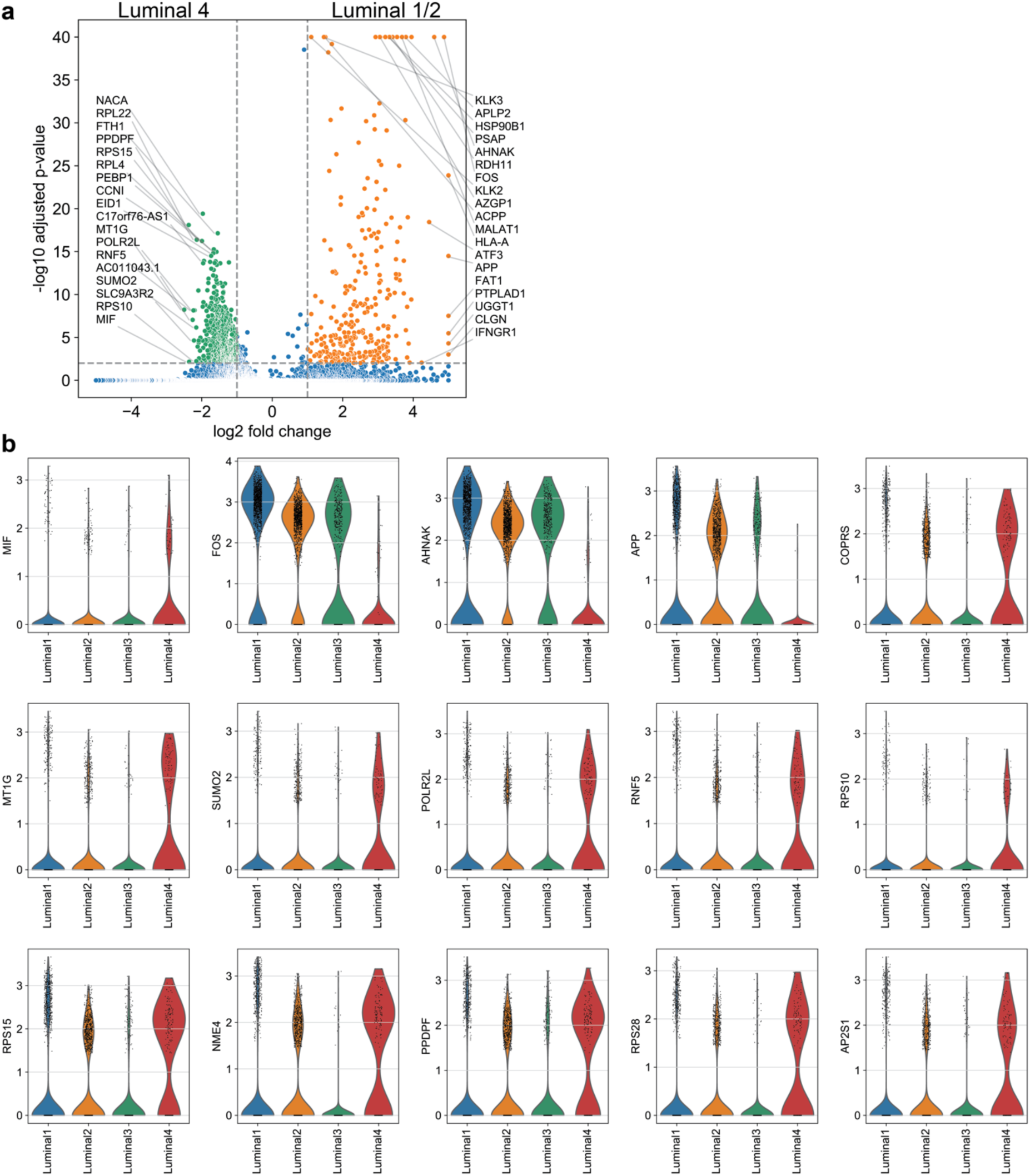
Luminal cell marker genes. (a) Volcano plot of differentially expressed genes found in the luminal 4 subgroup (green) or combined luminal 1 and 2 subgroups (orange). (b) Violin plots of marker gene expression in the 4 luminal subgroups.

**Supplemental Data 5.**
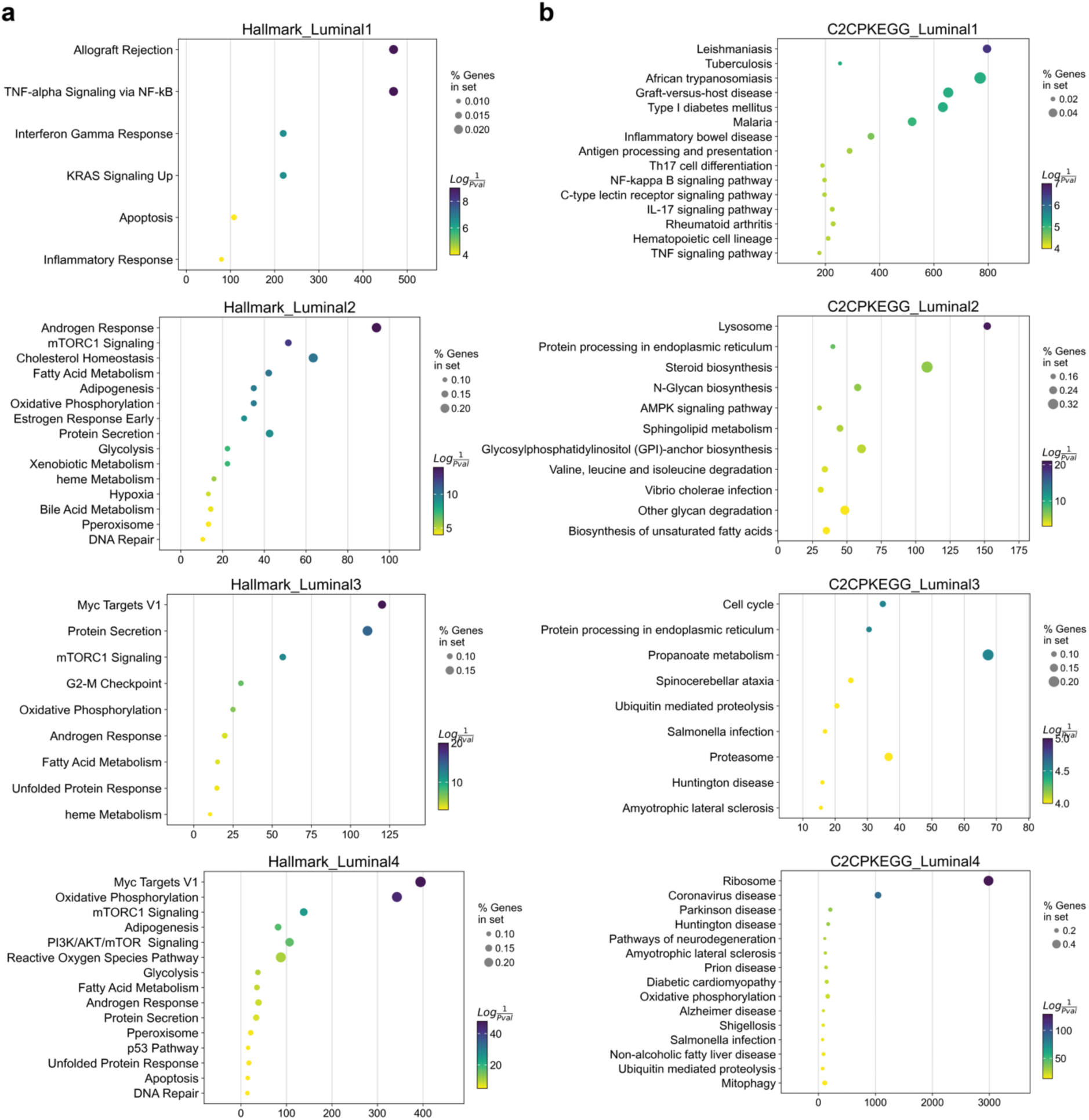
Gene set enrichment analysis of luminal subgroups. (a) Gene set enrichment analysis of differentially expressed genes from the 4 luminal subgroups. Genes were compared to Hallmark gene sets. (b) Gene set enrichment analysis of differentially expressed genes from the luminal subgroups. Genes were compared to gene sets from canonical pathways (CP) and Kyoto Encyclopedia of Genes and Genomes (KEGG).

**Supplemental Data 6.**
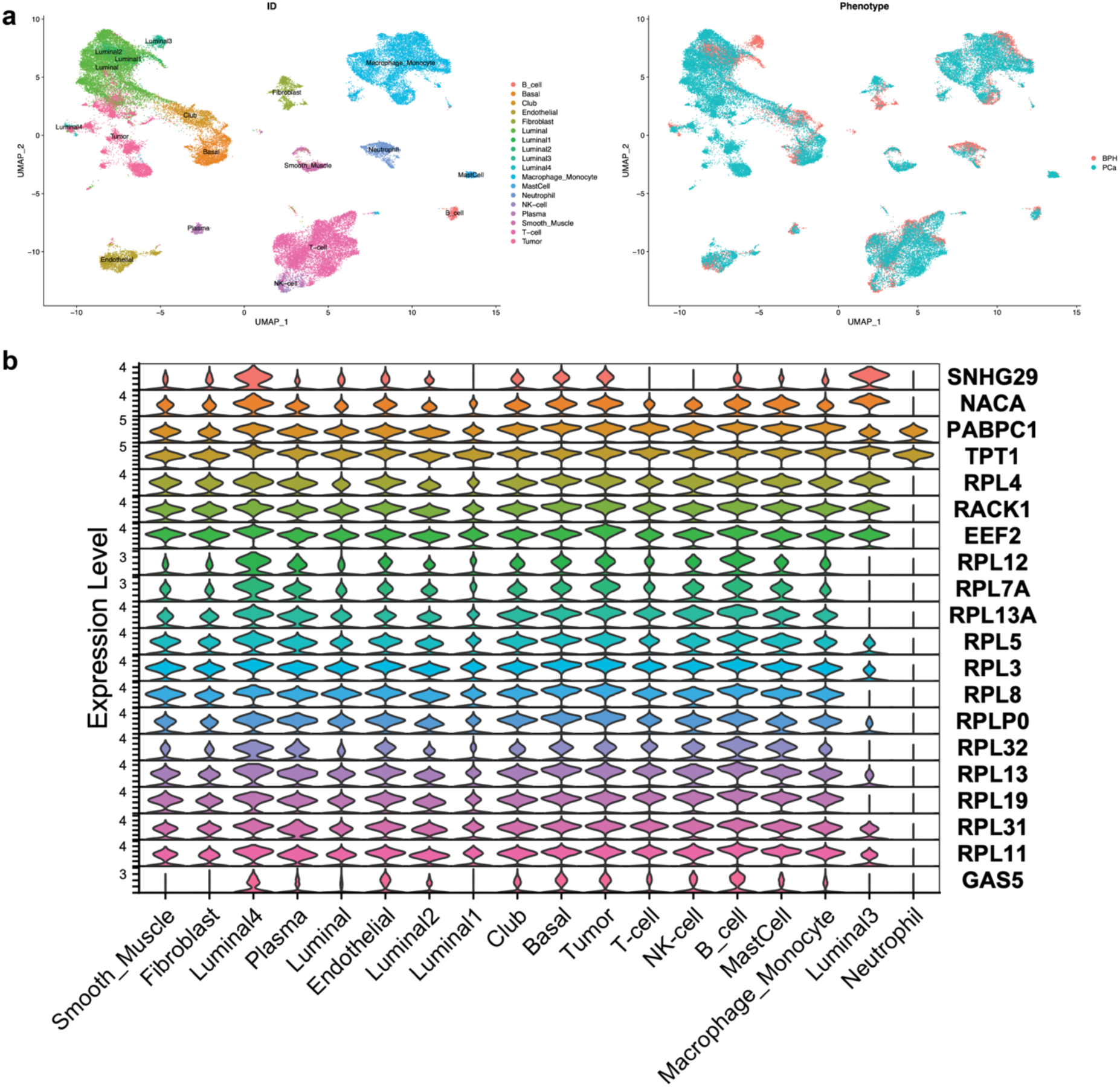
Detection of the luminal 4 subgroup in a prostate cancer dataset. (a) UMAP shows clustering of the integrated BPH and prostate cancer datasets. Cells were grouped by either by cell annotation or dataset. (b) Gene expression of the luminal 4 marker genes in the integrated dataset.

**Supplemental Data 7.**
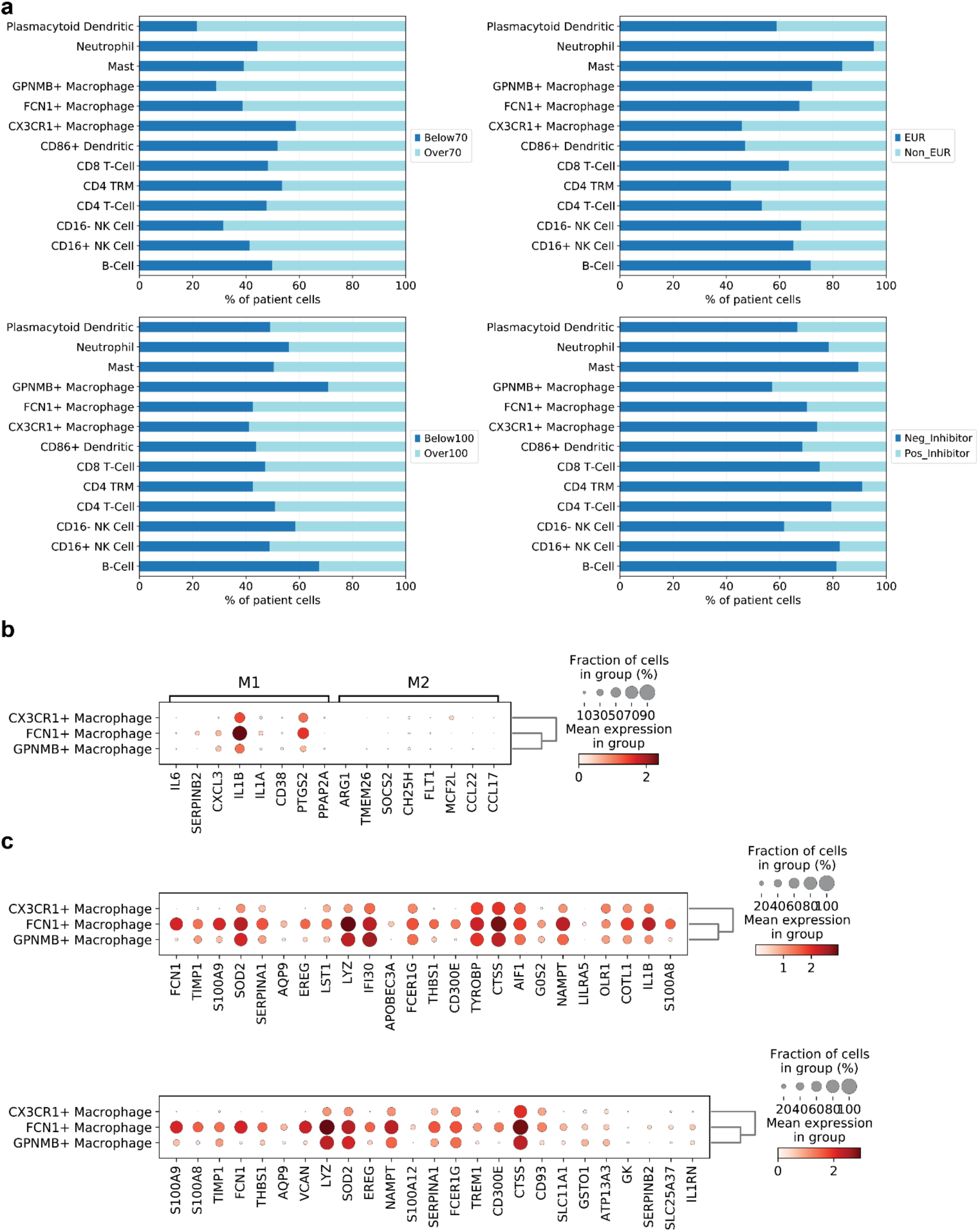
Immune cell characterization. (a) Cell contribution to immune cell groups from patients, split by age, size, ethnicity, and 5-ARI inhibitor status. (b) Dot plot of M1 and M2 macrophage marker gene expression in the 3 macrophage subgroups. (c) Dot plot of nonclassical (above) and classical (below) macrophage marker gene expression in the 3 macrophage subgroups.

**Supplemental Data 8.**
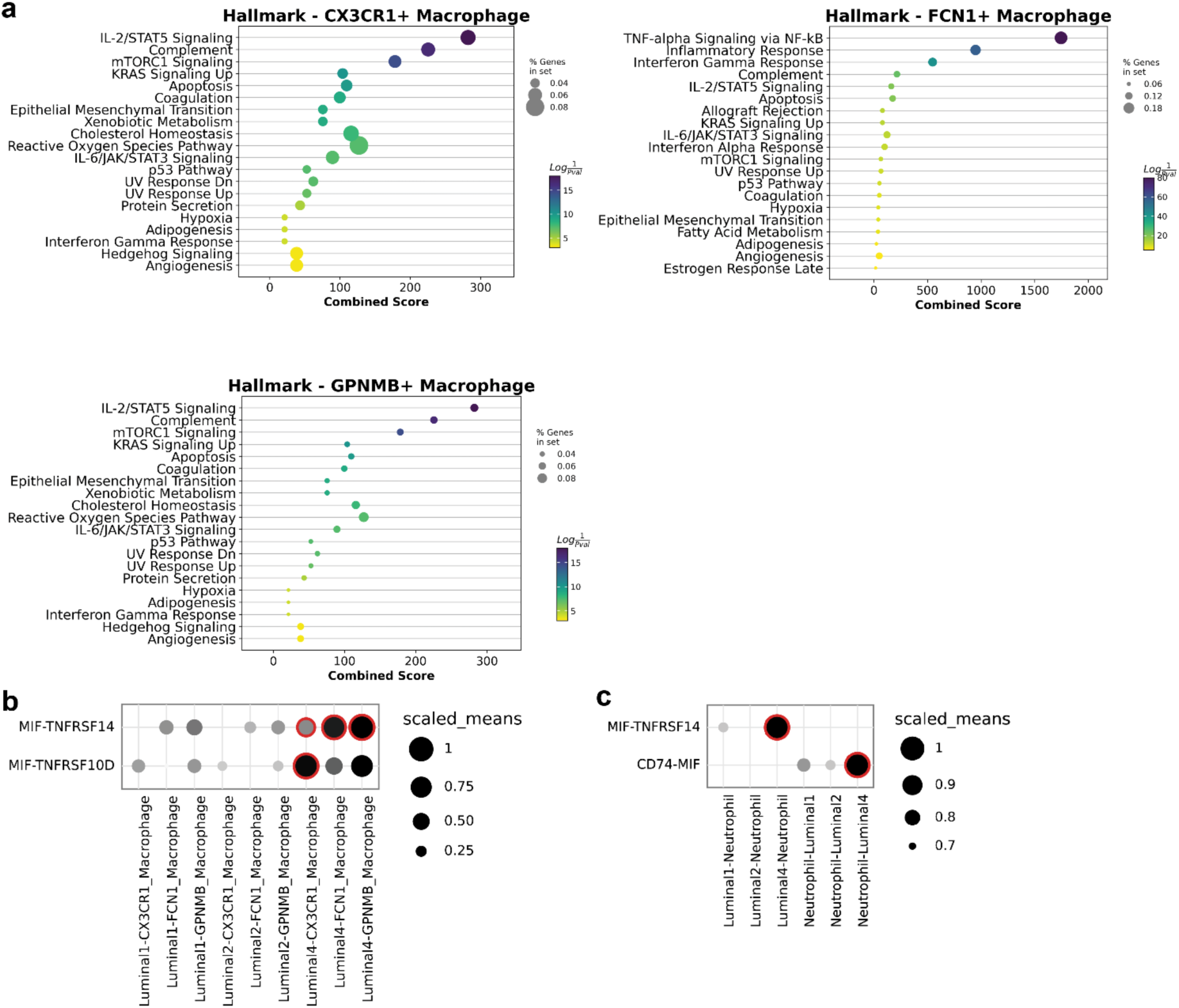
Macrophage gene set analysis. (a) Gene set enrichment analysis of differentially expressed genes from the CX3CR1+, FNC1+, and GPNMB+ macrophage subgroups. Genes were compared to Hallmark gene sets. (b) Interaction analysis between luminal subgroups and macrophages. Red highlights represent a significant interaction, with an adjusted p-value <0.05. (c) Interaction analysis between luminal subgroups and neutrophils. Red highlights represent a significant interaction, with an adjusted p-value <0.05.

